# Interventional Nerve Visualization via the Intrinsic Anisotropic Optical Properties of the Nerves

**DOI:** 10.1101/017046

**Authors:** Kenneth W.T.K. Chin, Andries Meijerink, Patrick T.K. Chin

## Abstract

We present an optical concept to visualize nerves during surgical interventions. The concept relies on the anisotropic optical properties of the nerves which allows for specific switching of the optical reflection by the nervous-tissue. Using a low magnification polarized imaging system we are able to visualize the on and off switching of the optical reflection of the nervous-tissue, enabling a non-invasive nerve specific real-time nerve visualization during surgery.

## 1. Introduction

Identification of nerves during surgical interventions is a key challenge, which is especially the case for the thinner nerves and nerves hidden by other tissue. It is of great importance to avoid accidental nerve injury, which can lead to decreased motion, loss of sensation or pain and even morbidity. [1] Several methods are currently available to localize the nerves in the body. For example MRI can be used to visualize nerves prior to surgery,[2,3] optical coherence tomography (OCT) and electromyographic monitoring are currently applied in delicate areas during surgery.[4,5,6,7,8] The limitations of these methods are that the nerves can only be visualized prior to surgery or do not give a specific, large area, and real-time nerve visualization. Currently, there is no clinical system available to visualize nerves in a safe, specific, and real-time way during surgery.[9]

In this work we present an optical concept to visualize nerves during surgery, using the anisotropic optical properties of the nerve cells. The extreme aspect ratio of the nerve cells compared to other cells in the body is a unique feature of nerves. This results in a strong anisotropic interaction with light. Oldenbourg et. al,[10] showed that the anisotropic optical behavior originates from oriented microtubules inside the nerve cells. Several spectroscopic studies showed small but distinct changes in scattering and optical polarization properties of nervous-tissue upon activation or electrical stimulation.[11,12] Backmann et al. describes that the polarized reflection of tissue is distributed over the solid angle ΔΩ and when the incident light is completely collimated (ΔΩ_0_= 0) the reflected polarized (backscattered) light will have a solid angle of ΔΩ= 0 parallel to the polarized incident light.[13] The anisotropic tissue reflection of the birefringent nerve tissue will therefore be most intense when the nervous tissue is aligned to the polarization of the collimated incident light. In our imaging concept this enables imaging of the anisotropic reflection of the nerves with a sufficient signal intensity at a relative long working distance (>10 cm), necessary in a surgical setting.

### Experimental

#### Animal tissue

All experiments on mice or chicken tissue were performed on tissue from as obtained dead animals. The adult dead mice were obtained from E-POS Globe Den Haag, the Netherlands.

#### Polarized light microscopy (PLM)

Nerves were imaged and validated using a Zeiss Photomicroscope I, equipped with III RS fluorescence filter block with 50:50 plate beam splitter obtained from Thorlabs and a Zeiss incident light side arm. Illumination was performed using a 100 W Zeiss/Osram halogen lamp. The incident light polarizer was obtained from Thorlabs. Image acquisition was performed using a Pixelink CCD camera attached to the external camera port of the microscope. Polarized transmission spectra were obtained using an Avantes 2048 CCD spectrometer. The fibre patch cord attached to the spectrometer included an integrated 3mm ball lens positioned in the focusing plane of the external camera port of the microscope.

#### Nervous-tissue validation

The validation of tissue samples to confirm the presence of nervous-tissue was performed using PLM. As discussed above the unique optical on/off switching of both the transmitted and reflected (red shifted) light upon rotation of the nervous-tissue enables the unique possibility to distinguish the nervous-tissue from other types of tissue. Furthermore the tissue specific reflection or transmission using PLM also enables the visualization of the tissue structure down to the cellular level, enabling the identification of individual nerve cells within the tissue sample.

#### Surgical imaging

An imaging setup as disclosed in Fig. 1a was used to perform surgical imaging on the mice, and chicken legs. The system comprises a Pixelink CCD camera equipped with a Fujian c-mount camera lens (f=35 mm), depending on the desired magnification one or two additional imaging lenses (f= 15.5 or 12 cm) where used. The imaging polarizer was obtained from Thorlabs and the illumination polarizer was a standard polarizing laminated film.

**Figure 1:**
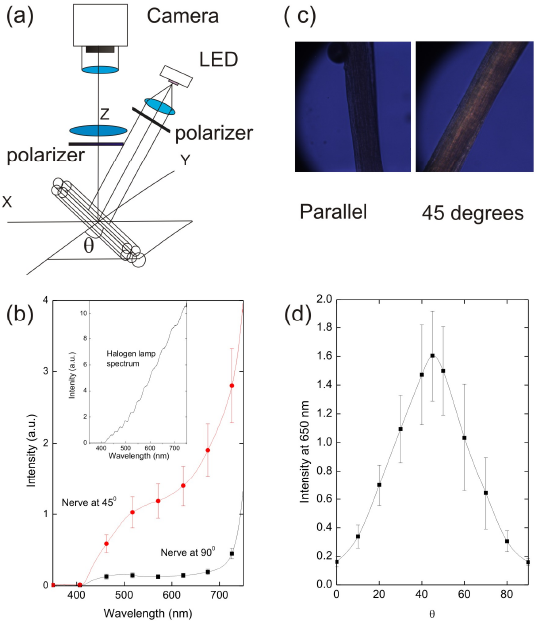
(a) Representation of the collimated polarized light imaging system with a nerve. (b) The anisotropic optical properties of thin slices of nervous-tissue where measured using PLM in transmission mode. The fraction of transmitted light through a nerve as function of the wavelength. (c) PLM image of a chicken nerve, (10× objective) parallel and at θ=45 •with respect to the incident light polarization. (d) Polarization dependence of the transmitted light intensity trough nervous tissue as function of the rotation angle (θ).

### Results and discussion

The easily accessible sciatic nerve from a mouse was chosen to demonstrate the anisotropic properties on which our imaging concept is based. The nerve imaging system (Fig. 1a) comprises a collimated polarized light source and a camera system to measure the reflected light with perpendicular polarization. To validate the anisotropic optical properties of nervous-tissue we first studied the sciatic nerve of a mouse with polarized light microscopy (PLM) in transmission. The sciatic nerve is myelinated, relatively thick and comprises a significant amount of connective tissue, reducing the anisotropic optical properties by scattering so thin slices of this nerve were used to reduce the isotropic scattering. The nerve tissue reveals an enhanced red shifted transmission intensity (Fig. 1b,c), when oriented to 45° degrees with respect to the cross-polarizers. After aligning the nervous-tissue with either one of the cross polarizers a reduced reflection (at •>500nm) of the nerve was observed (Fig 1b,c, parallel). Fig. 1d confirms that the measured polarized tissue transmission at • 650 nm maximizes at 45° with respect to either the incident light- or imaging-polarizer, which is consistent with previous spectroscopic findings for different nerve types,[11,12,14] confirming the potential for specific nerve visualization by their anisotropic properties.

The ability to visualize nerves in the presence of other tissue types is very important during surgery. Fig. 2a shows the ability to distinguish between nerves embedded in the fat in a section of chicken skin, using PLM by their specific orange appearance upon rotation under polarized light. The nerves (indicated by the green arrows) in the fat tissue appear with an orange signature at 45° in contrast to the isotropic transmission of the fat cells, appearing grey. Muscle-cells also possess an anisotropic optical behavior as a result of their cellular composition and shape. Nevertheless using PLM nervous-tissue can be distinguished from the muscle-tissue by their selective appearance upon rotation with respect to the polarized incident light as shown in Fig. 2b showing a nerve (indicated by green arrows) embedded in a muscle (dark texture).

**Figure 2.**
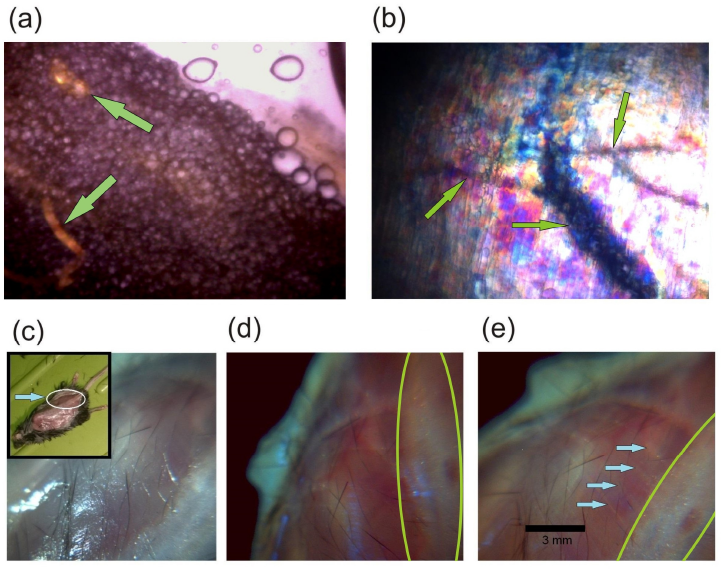
(a) PLM image of nerves (orange) indicated by the green arrows in fat tissue (chicken), (b) PLM image of a nerve seen as a black shadow (indicated by the green arrows) embedded in muscle tissue (mouse), this tissue section was dissected from the same area as indicated in Fig. 2c-e. (c) Surgical setting where the skin at the back of a mouse is opened showing the muscles and the spine under incident white light, the inset shows the mouse with the AOI. (d) cPLI image with the spinal cord perpendicular to the imaging polarization. (e) Rotating the mouse with the spinal cord oriented to 45° with respect to the cross polarizers shows the visual (orange, marked by the blue arrows) appearance of nerve-roots (• 0.05 mm in diameter) originating from the spinal-cord (indicated in the area surrounded by the green line).

A microscopic setting is ideal to visualize the anisotropic tissue properties in a small area at a relative short distance with respected to the imaging objective lens. However, during surgery polarized light microscopy is not easy to use due to the relative large size of the area of interest (AOI), tissue diversity, and surface topology. Polarized light imaging has been used to study tissue fiber orientations in thin brain tissue slice.[15] However, with “normal” non-collimated polarized light it is not possible to image specifically the nerves in thick tissue as encountered in a surgical setting. For imaging in a surgical setting we developed a collimated polarized light imaging (cPLI) system depicted in Fig. 1a, comprising a collimated polarized light source and a camera for detecting the reflected polarized light. Using this system we imaged the back of a mouse with the skin removed (Fig. 2c, the inset clarifies the AOI). With non-polarized illumination it is impossible to indicate if and where the nerve-roots originate from the spinal cord. However, using cPLI a distinct appearance (Fig. 2e, orange lines) and disappearance (Fig. 2d) of the nerve-roots embedded in muscle tissue is observed when rotating the mouse with respect to the polarization direction. The visualized diameter (• 50 μm) of the nerve-roots is about one order of magnitude smaller than the surgically most relevant nerves.

Nerves in chicken legs were used to evaluate the cPLI system in a size range closer to the human situation (~0.5 mm). Depending on the surrounding tissue of the nerves, the incident light scatters isotropically reducing the orange specific reflection observed using PLM. However, Fig. 3a,b shows that nervous-tissue can be distinguished from the surrounding tissue by their *on* and *off* switching when rotating the chicken leg with respect to the polarization direction of the cross-polarizers. Parallel to the incident light the nerve (situated at the tip of the needle) appears transparent, and rotated to • 45 ° the nerve reveals a stronger white reflection. Fig. 3C shows the tissue validation by PLM, revealing the nervous-tissue structure and the specific orientation dependent anisotropic reflection, used for confirmation of the presence of nervous-tissue. Under non-polarized illumination it is often difficult to identify nerves during surgical interventions. Fig. 3d shows an opened chicken leg under normal white light illumination. It is impossible to indicate which of the tissue patterns corresponds to the nervous-tissue. However, by rotating the sample under collimated polarized light the narrow tissue string (blue arrows) almost disappears at 90° degrees and reappears at +/-45° degrees indicating the presence of nervous-tissue. Furthermore, the nerve splits into two branches (black and green arrows), which are not visible under normal illumination.

**Figure 3.**
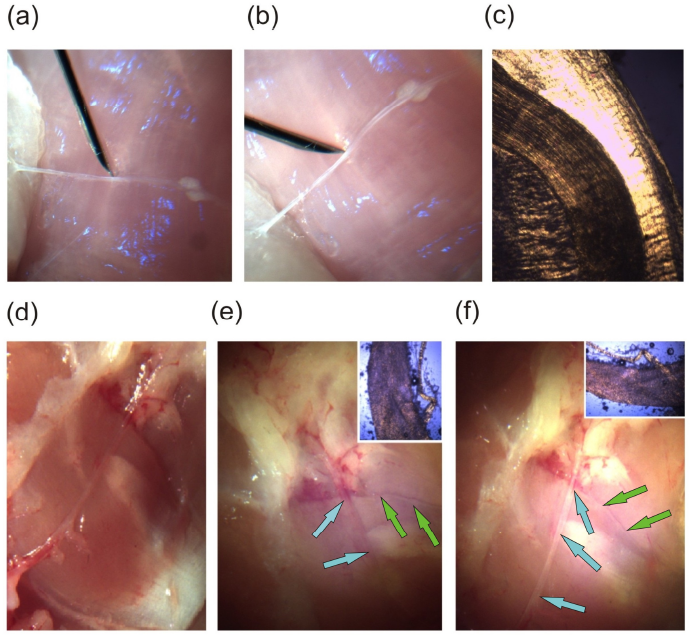
cPLI of a nerve in a chicken leg, (a) showing a nerve (diameter • 0.3 mm) parallel to either one of the cross-polarizers. (b) Upon rotation of the nerve at 45° the nervous-tissue appears white. (c) Validation of the nerve tissue by PLM. (d) Chicken tissue under normal white light, (e) rotating the sample using cPLI shows the almost disappearance of the nervous-tissue (diameter<0.4 mm, blue arrows). (f) Rotating this structure to 45° with respect to the cross-polarizers it appears white. The insets show the validation of the nervous tissue by PLM.

Due to the absorbing and scattering properties of myelin and connective-tissue it was not possible to identify “thicker” nerves (>1 mm in diameter) using cPLI. The additional presence of myelin and connective-tissue depolarizes and scatters the incident light significantly reducing the visibility of the nerve. Fortunately, these thicker nerves are often easily recognized by the surgeon without the need of an imaging technique.

### Conclusion

Using the unique anisotropic optical properties of the nerves we were able to visualize nerves up to only a few nerve cells buried in fat. By applying a collimated polarized light source combined with a perpendicular oriented polarized camera it is possible to recognize thin nerves in a surgical setting in mice and chicken legs. The selective appearance and disappearance of the of the thin (< 0.5 mm in diameter) nerves upon rotation with respect to the optical polarization enables real time, specific, and safe identification of nerves during surgical interventions.

